# Camouflaging of repetitive movements in autistic female and transgender adults

**DOI:** 10.1101/412619

**Authors:** Joost Wiskerke, Heléne Stern, Kajsa Igelström

## Abstract

Repetitive movements (RMs), colloquially called “stimming” among adult autistic people and “motor stereotypies” among scientists, are common in autism. These behaviors fall under the domain of restricted and repetitive behaviors in the current edition of the Diagnostic and Statistical Manual of Mental Disorders (DSM-5). RMs can be socially disruptive or cause self-harm, but can also be experienced as cognitively or emotionally helpful and even enjoyable. Overt RMs are less common in females than in males, which could contribute to clinical difficulties in detecting their autism. In the social domain, autistic people with intact intelligence can often mask their social difficulties through various compensation strategies, and females appear especially skilled at it. Subjective report from verbally able adults may be useful as a first step in detecting potential camouflaging of RMs, and to provide a foundation for further studies. We founded an Internet-based outreach platform that became particularly successful in reaching female and transgender individuals. We recruited 342 individuals to an anonymous online questionnaire, collected data about self-reported RMs and probed for potential camouflaging. The cohort comprised 56% formally diagnosed participants and 44% who self-identified as autistic, and 17% of all participants reported non-cisgender identity. Thus, in addition to diagnosed women, we reached two populations that would normally be excluded from autism studies: transgender and undiagnosed participants. We found high rates of RMs in both diagnosed and self-identifying participants, and a striking prevalence of camouflaging. We suggest that camouflaging of RMs may contribute to underdiagnosis of autism, at least in females and transgender people, and that further studies on this topic are exceptionally important.

## Introduction

Awareness of the behavioral and neural diversity of the autism spectrum is on the rise, and barriers to early detection and diagnosis are actively discussed by researchers, clinicians and society as a whole (Lai and Baron-Cohen 2015, Lewis 2017, Crane *et al.* 2018). One difficult challenge in recognizing autism in cognitively able individuals is that it is common to “camouflage” difficulties in response to social demands (Kopp and Gillberg 1992, Kopp and Gillberg 2011, Hull *et al.* 2017). Camouflaging in the social domain has been recognized and studied in the context of the underdiagnosis of autism in girls and women (e.g. Bargiela *et al.* 2016, Lai *et al.* 2016). The possibility of a “female phenotype” has been discussed, although we should not forget the possibility that some males too may camouflage and escape diagnosis in a similar way (Kopp and Gillberg 2011, Bargiela et al. 2016). Autism is especially underdiagnosed in adults – a population that has been referred to as “the lost generation” (Lai and Baron-Cohen 2015) and whose accumulated comorbidities often complicate diagnostic assessment (Russell *et al.* 2016).

Autistic children often express rhythmic non-functional repetitive movements (RMs), which may or may not persist into adulthood. They differ from tics in that they do not begin with an urge to move and are often oscillatory and more prolonged in nature (Mills and Hedderly 2014). RMs occur in typical development and are not specific for autism – anyone can click their pen, bounce a leg, or bite their nails. However, RMs are more common in autistic than in typically developing children, and have been suggested as a target for intervention, in part because they have been correlated with the severity of other deficits and in part because they might interfere with other functions (Matson and Dempsey 2008, Lanovaz *et al.* 2013). For example, some concerns among clinicians are social stigma, self-injury and missed educational opportunities (Loftin *et al.* 2008). However, autistic individuals do not necessarily mind their RMs and may report them as pleasurable or helpful (Singer 2013).

RMs are only one possible expression within the heterogeneous DSM domain of restrictive and repetitive behaviors (RRBs), which includes behaviors ranging from simple stereotypies to higher cognitive RRBs such as circumscribed interests or ritualistic behaviors (and, since DSM-5, also atypical sensory function). Thus, in the clinic there is often no need to characterize RMs in depth, as long as enough RRBs are present to establish a diagnosis. In contrast to higher-level RRBs, RMs do not require intact cognitive functions. They decrease with social maturation and age, and are more common in individuals with intellectual disability (ID) (Loftin et al. 2008, Esbensen *et al.* 2009). Research studies of RRBs in autistic individuals without ID have naturally focused more on higher-order RRBs and sensory issues than lower-order RMs (South *et al.* 2005). For these reasons, and because RMs are quite harmless in many cases, we do not know much about the prevalence of RMs in adults without significant ID. Given the ability of many autistic women to hide their social deficits (Hull et al. 2017), there is no logical barrier to the hypothesis that RMs could occur together with intact cognitive function and conscious self-reflection, and be camouflaged in some way. Increased knowledge about RM expression and possible camouflaging may help both in clinical assessment and in studies of neural correlates of RRBs and their potential function. In addition, a better understanding of the autism spectrum may ultimately lead to better societal acceptance and improved well-being of autistic individuals.

In this exploratory study, we took a first step in testing this hypothesis by using self-report measures from a large number of female and transgender adults, using recruitment on social media to an online questionnaire. Throughout this report, we use identity-first phrasing and non-pathologizing language, as this is the preference of the majority of our participants (e.g. “autistic person” rather than “person with autism spectrum disorder”) (Kenny *et al.* 2016, Gernsbacher 2017).

## Methods

The study was approved by the Princeton University Institutional Review Board. We recruited participants on Facebook, using short advertisements that contained a link to an external website where more information was available. The anonymous questionnaire was run on the Qualtrics platform, without using cookies or saving IP addresses. Participants were not reimbursed for their participation.

### Recruitment and participants

In designing the study, we built on our experience from a series of exploratory questionnaires run during 2017, which had resulted in extensive constructive feedback from the community. In preliminary unpublished studies, recruitment was found to be most successful when we were able to collaborate with administrators of closed, well-managed groups with 1000–2000 members. Ads posted within groups and on our own page were spread further through sharing. In addition, a self-advocacy group advertised the study, and we used Twitter as another (but for us less successful) venue.

We recruited female and transgender/non-binary adults who identified as autistic and/or had a formal diagnosis of an autism spectrum condition. We did not allow cisgender males (men who were assigned male at birth and also identified as male) to participate, because we had been unable to recruit meaningful numbers of cisgender men to preliminary studies. We required honesty and disclosure of diagnostic status and gender identity, but assured the participants that the information would not be used to exclude anyone.

### Terminology decisions

The online questionnaire was designed, based on feedback in preliminary studies, to be autism-friendly (e.g. minimal ambiguity and carefully phrased response options) and to minimize negative reactions, unpredictability and cognitive overload. We added textboxes on every page, which helped participants get through sections that they struggled with. The most common reason for opting to use those was when a precise-enough match could not be found in multiple choice options. Being able to add disclaimers and explanations helped participants complete the questionnaire and gave us information about any problematic sections. Most questions were optional, to allow participants with limited energy to participate despite the need to complete the questionnaire within one browser session.

Another barrier we had previously encountered was that a proportion of participants reacted very negatively to language that they interpreted as stigmatizing or pathologizing, making it particularly difficult to use standardized self-rating measures, such as the autism quotient (AQ) (Baron-Cohen *et al.* 2001). For this study, it was particularly important to be able to use words such as “symptoms”, and to have an AQ score for comparison with other populations in the literature. Therefore, we started the questionnaire with a note on terminology, which contained an acknowledgment of the complexity and importance of the debate surrounding person-first versus identity-first language, and that our decision to use identity-first language was based on what seemed to be the most common preference. We also acknowledged that some of the questions may sound clinical or pathologizing, but that it was sometimes necessary for us to phrase questions in such a way. We also gave a short overview of the questionnaire, which consisted of 8 pages of different lengths.

### Addressing potential biases

We decided to inform, and keep reminding, the participants about the possibility that they may not identify with any of the behaviors mentioned and that we were not testing whether or not they were likely to be autistic. The main reason for emphasizing this was that we had seen that many people in adult populations had a very painful history, involving many years of feeling unseen and disrespected, even after a formal diagnosis had been made. To avoid any unconscious biases based on common perceptions of what autism should “look like”, we wanted them to be onboard with us in exploring their own experiences as unchartered territory.

Trial and error in preliminary studies had shown that the highest quality data and best participant satisfaction were obtained by 1) starting with some multiple-choice options to avoid the need to organize a response completely from scratch, and 2) following with an optional textbox for elaboration. This resulted in many detailed text responses and high response rates. However, we stayed as neutral as possible, minimizing order bias in multiple-choice questions by randomization, and using open-ended phrasing in textbox questions.

### Demographic and clinical questions

We asked about age, country of residence, employment status, level of education, sex assigned at birth, and gender identity. After consultation with a representative of the transgender community, we chose to ask about “sex assigned at birth” (“Female”, “Male” or “Other”), as well as about “gender identity” (“Female”, “Male”, “Unsure”, or “Other/no answer”, with a textbox connected to the latter).

On a new page, we asked the following question: *“Have you been diagnosed with Autism Spectrum Disorder, Asperger Syndrome, PDD-NOS or other autism spectrum condition?* Our goal with this question was to divide participants into diagnosed and self-identifying groups.

Participants with a formal autism diagnosis were asked what type of health professional gave them the diagnosis. Options included psychologist, neuropsychologist, psychiatrist, neurologist, speech therapist, pediatrician, general practitioner/primary care physician, and “other (please specify)”. An extra textbox was provided for elaboration or clarification. We also asked about year of diagnosis and age at diagnosis, and finished with a general textbox for comments.

We also asked participants *“Have you ever been diagnosed with any of the following? Please include diagnoses that you think are misdiagnoses.”* The options included other psychiatric and neurodevelopmental diagnoses, including attention deficit/hyperactivity disorder (ADHD), dyslexia, developmental coordination disorder (DCD; dyspraxia, apraxia), ID, anxiety disorder, depression, eating disorder (with option to specify), obsessive/compulsive disorder (OCD), post-traumatic stress disorder (PTSD), bipolar disorder, schizophrenia, personality disorder (with option to specify), and other conditions (with option to specify). At the top of the list, the options “I have no conditions except autism” and “I prefer not to answer this question” were presented, followed by the conditions in randomized order, and we listed the option “other conditions” at the bottom of the page. We then provided a textbox with the instruction *“Please add any comments you might have about the previous question. If you think you have been misdiagnosed with any of the conditions above, please mention it here.”*.

Given the non-specific and subjective nature of our question (including all diagnoses ever received), these data were not used for statistical comparisons across groups. However, they provided a general view of the clinical history of our participants and the profiles within the two groups.

### Assessment of RM expression and camouflaging

We assessed current RMs using a combination of visual analog scales for specific behaviors and textboxes for free-text responses. For the self-rating question, we used examples of stereotyped or repetitive motor movements based on criterion B1 in the DSM-5 (stereotyped or repetitive motor movements, use of objects, or speech). We did not include speech (e.g. echolalia, idiosyncratic phrases) as we thought this might need in-person assessment to a greater extent than the other behaviors. However, we did include one option for “repeating certain sounds”. The scale ranged from “Never” (0) to “Very often” (10) and had a resolution of two decimal points.

The instructional text was designed to span the terminology used in the autistic community, and express that motor stereotypies or self-stimulatory behaviors are not in and of themselves necessary for an autism diagnosis (only two out of four of the DSM-5 RRB examples must be fulfilled). The following text was used:

*“On this page, we probe to what extent autistic women have what clinicians call “motor stereotypes” or “repetitive movements”. Autistic individuals often call it “stimming”, so we use that term too for simplicity. Not all autistic people do stimming. If you are not a person who stims, please just leave the slider bars at zero. If you think you express stereotyped or repetitive movements more than most neurotypical people, and/or consider yourself someone who stims, please use the slider bars below to rate how often you’ve engaged in the following behaviors in the past 3 months, on a scale ranging from “Never” (0) to “Very often” (10). We understand that it’s not possible to give a precise answer.”* Next, we asked (free-text boxes): *“Did you engage in any of the behaviors above when you were younger? If so, which ones and approximately at what age?”*, and *“Do you, or did you ever, have other stimming patterns/repetitive movements that are not mentioned above? Please describe in as much detail as you like.”*.

Camouflaging was probed using the matrix multiple-choice question *“Did/do you hide these behaviors from others…”* 1) *“…as a child?”,* 2) *“…as an adolescent?”,* 3) *“…as an adult?”.* The options were “Always”, “Sometimes”, “Never”, “Don’t know”, and “N/A”. After this, we asked *“If applicable, please tell us about how you have been and/or are hiding stimming behaviors.”*. Lastly, we asked *“Do you wish you could control these behaviors better?”* with the multiple-choice options “N/A”, “Yes” and “No” – the latter two with optional textboxes in case of comments. When we asked about the life phases of childhood, adolescence and adulthood, we defined those as ≤12 years, 13–17 years, and ≥18 years, respectively.

### The AQ

On the last page, we presented the AQ, and made it mandatory. We knew from preliminary studies that many of our participants struggle with and/or dislike this questionnaire, so we designed our introductory text to acknowledge this: *“Finally, we ask you to please fill out the following questionnaire. It is a test used by many other researchers, which means that we need it for comparison purposes – we do not use it to “test” your autism, because it might be male-biased. Please respond to all the items to the best of your ability even though we know this page can be difficult.”*. We provided a textbox at the bottom of the page to let the participant explain their choices or give other feedback. This approach was well-received, except for one diagnosed participant who filled it out randomly “in protest”.

### Analysis

Statistical tests were performed in Prism 7 (GraphPad Software Inc., CA, United States), were non-parametric in all cases, and used Bonferroni correction for multiple comparisons when appropriate. Text responses were analyzed using Quirkos 1.5.1 (Quirkos Ltd, Edinburgh, Scotland) and MATLAB R2018a for cluster analysis (MathWorks, Inc., MA, United States). We avoided analyses that might mislead a reader to generalize results beyond the demographic groups that participated in our study, and reported these as qualitative results instead.

## Results

We recruited 342 cis-gender female and transgender individuals (18–80 years old) who identified as autistic, with or without a formal diagnosis. The main reasons for choosing a fully inclusive study were that it 1) would enable valuable communication with “self-identifying” individuals, who are excluded from conventional autism studies, and 2) may encourage participants to be honest about their diagnostic status as there were no negative consequences to lacking a diagnosis.

### Participants

Most participants in both groups lived in the USA or UK (**Fig. 1A**), and 92% of all participants lived in countries with English as an official language (USA, UK, Ireland, Canada, Australia, and New Zealand). However, the study also attracted some participants in various other countries (Sweden, Finland, Denmark, Norway, Iceland, the Netherlands, Germany, France, Spain, Hungary and Turkey).

**Figure 1.**
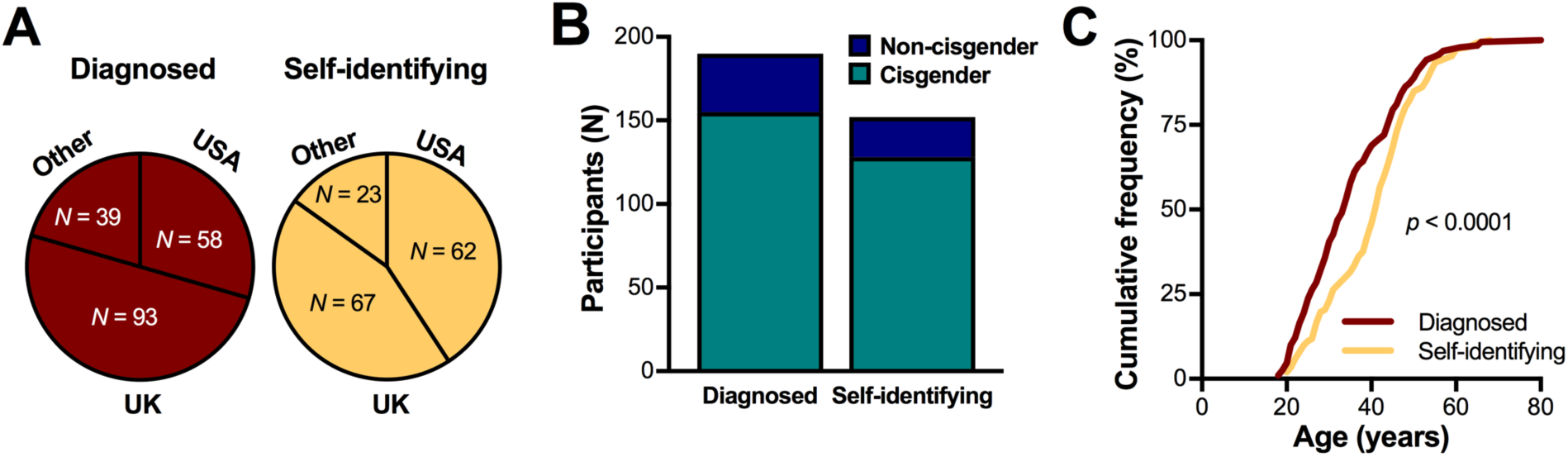
Demographic data of the diagnosed and self-identifying groups. A. Participants in both groups were predominantly located in the United States of America or the United Kingdom, with a smaller proportion of individuals living in other countries (see Participants section). B. Overview of the number of participants in the two groups, broken down into cisgender females and non-cisgender individuals. C. The cumulative frequency of ages across diagnosed and self-identifying groups, showing a greater proportion at younger ages in the diagnosed group (*p* < 0.0001).

The sample included 190 participants (35 non-cisgender) with a formal autism diagnosis, and 152 participants (24 non-cisgender) who self-identified as autistic (**Fig. 1B**). We pooled cisgender and non-cisgender individuals, because our gender identity question was not detailed enough to enable reliable comparisons between cisgender females and the very diverse non-cisgender group. Of the participants who did not choose “female” as their gender identity, 55 had been assigned female at birth and 4 had been assigned male. Twenty individuals indicated that they were unsure about their gender identity, 26 provided definitions that were not binary (e.g. non-binary, gender-fluid, gender-queer, agender, demigirl), one was a male assigned female at birth, and three were females assigned male at birth. Nine participants provided unclear answers, either by leaving the textbox blank (7 participants) or by leaving comments along the lines of not agreeing with the concept of gender.

The age distribution of the diagnosed group showed a somewhat younger profile than that of the self-identifying group (median ages 34 and 41 years, respectively; *p* < 0.0001, Kolmogorov-Smirnov test; **Fig. 1C**). Within both groups, 99% reported having finished high school, and 91% in the diagnosed group and 93% in the self-identifying group had proceeded into college, indicating that the vast majority were indeed very cognitively able. In the diagnosed group, 56% reported that they were either working or studying, 16% that they were currently disabled, 8% unemployed looking for work, and 3% retired. The remaining 17% indicated that they were unemployed not looking for work. In the self-identifying group, the corresponding numbers were 66% working/studying, 8% disabled, 7% looking for work, 4% retired and 16% not looking for work. Several participants expressed dissatisfaction that they as stay-at-home mothers had to mark “unemployed not looking for work”, which should be taken into account when considering this status.

All participants above 25 years of age (76% of the diagnosed group), with one exception, had been diagnosed after the age of 15. The age of diagnosis is plotted against current age in **Figure 2A**, illustrating that the majority of participants were relatively newly diagnosed (Spearman r = 0.906). 65% of the group had received a formal diagnosis within the past 5 years and 82% in the past 10 years (**Fig. 2B**). The distribution of AQ scores (**Fig. 2C**) was similar to those of autistic participants in early studies (e.g. Baron-Cohen et al. 2001), and there was no significant difference between diagnosed and self-identifying participants (Kolmogorov-Smirnov test; *p* = 0.092).

**Figure 2.**
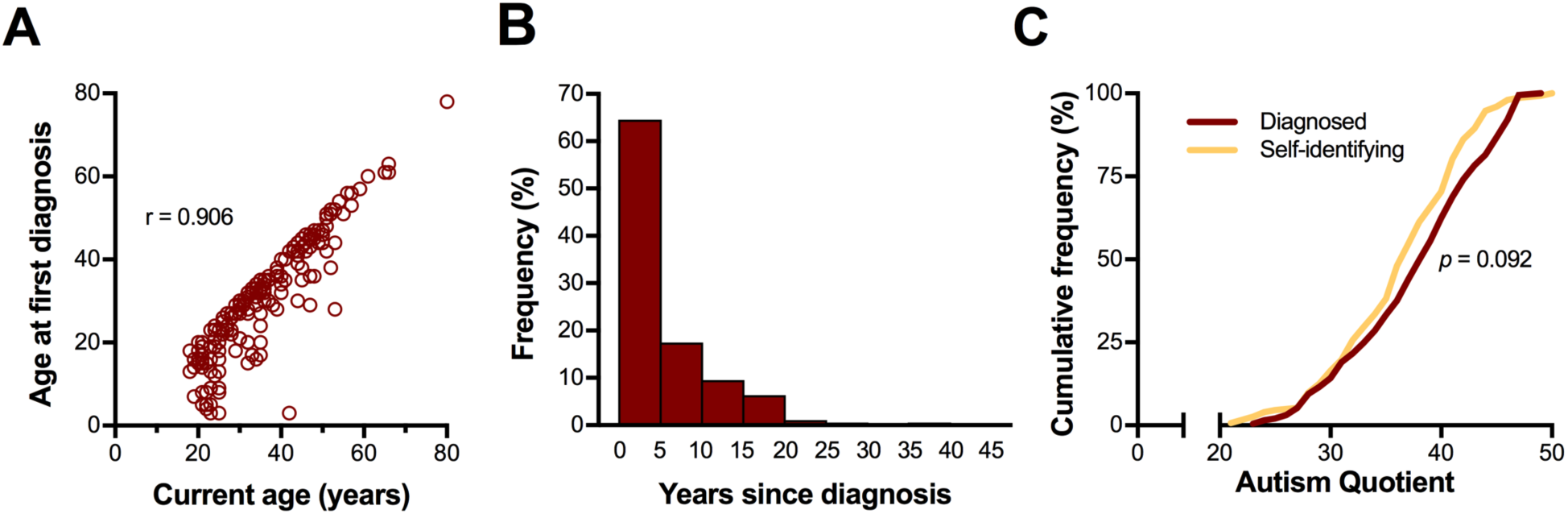
Age at diagnosis and AQ scores. A. Visualization of how the age at diagnosis compared to current age in all participants, showing that diagnosis in childhood was almost exclusively seen in the youngest participants and that the majority had been diagnosed very recently. B. Quantification of years since diagnosis, showing that the majority had received a diagnosis within the last five years. C. The AQ score distribution did not differ significantly between diagnosed and self-identifying participants.

### Comorbidities

The study format was not suited to determining true comorbidity rates (sampling bias, geographical diversity, and potential subjectivity). However, we asked participants to mark on a list of psychiatric and neurodevelopmental conditions or write in a textbox any diagnoses they had ever received, including those that had been removed or that participants considered to be incorrect. A textbox was included for any “other conditions”, and there was space to elaborate on personal opinions about the diagnoses.

One participant checked the “prefer not to answer” box, and specified that she could not cope with the negative emotions involved in going back through her records after having been misdiagnosed many times before her autism diagnosis. All other participants opted to respond.

Qualitatively, the group profiles were similar (**Fig. 3**). ADHD was reported by almost a fifth of participants in both groups, whereas the neurodevelopmental conditions dyslexia and DCD were reported more commonly in the diagnosed group in this cohort. Depression and anxiety were most common (>57% in both groups) and frequently comorbid with each other (45% and 52% of diagnosed and self-identifying participants, respectively). An eating disorder was reported by around 15%, with anorexia, bulimia and binge eating disorder being specified most frequently (3–6% each; similar across groups). OCD was reported by >12% in both groups, and PTSD was reported by around a fifth of participants (**Fig. 3**).

**Figure 3.**
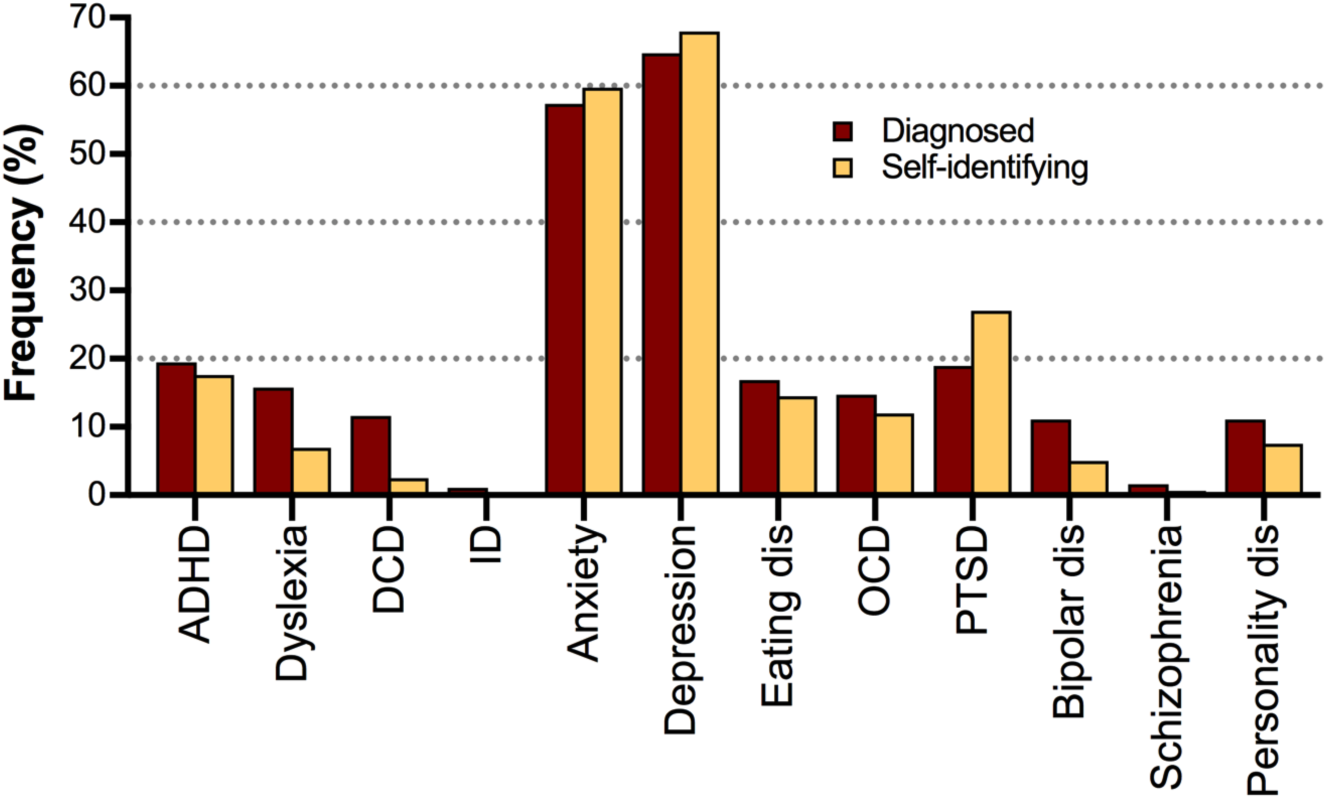
Other diagnoses ever received. Shown are the percentage of each group who reported having received the listed neurodevelopmental and psychiatric conditions at some point in their lives, regardless of whether they reflected misdiagnoses.

Overall, it was common with comments about misdiagnoses due to undiagnosed autism (especially bipolar disorder and borderline personality disorder), doubts about certain diagnoses (commonly personality disorders), and additional conditions suspected by the participant (in particular ADHD and DCD). One senior participant also mentioned that she had received a schizophrenia diagnosis at a time when autism still fell under the schizophrenia umbrella.

### Non-functional movements or use of objects

We first presented sliding bars for 12 RMs, including “repeating certain sounds”, and asked participants to rate how often they had engaged in each behavior *in the past 3 months*. The range of the analog scale was 0 (“Never”) to 10 (“Very often”). A few participants (6%) provided a disclaimer of having limited awareness of their RM, such as having had to ask family members for feedback or that they sometimes would not notice unless someone commented on it. Apart from that, there were no indications that participants had trouble answering this question. The interquartile ranges of self-ratings for each listed behavior are shown in **Figure 4**, with the different behaviors sorted in descending order based on the median score in the diagnosed group. With the exception of head banging in the diagnosed group, all behaviors were rated as 10 out of 10 by at least a proportion of participants, and were reported to at least some degree by >25% of participants. Hierarchical cluster analysis did not reveal any subgroups with specific patterns of RMs; rather, the patterns seemed extremely variable.

**Figure 4.**
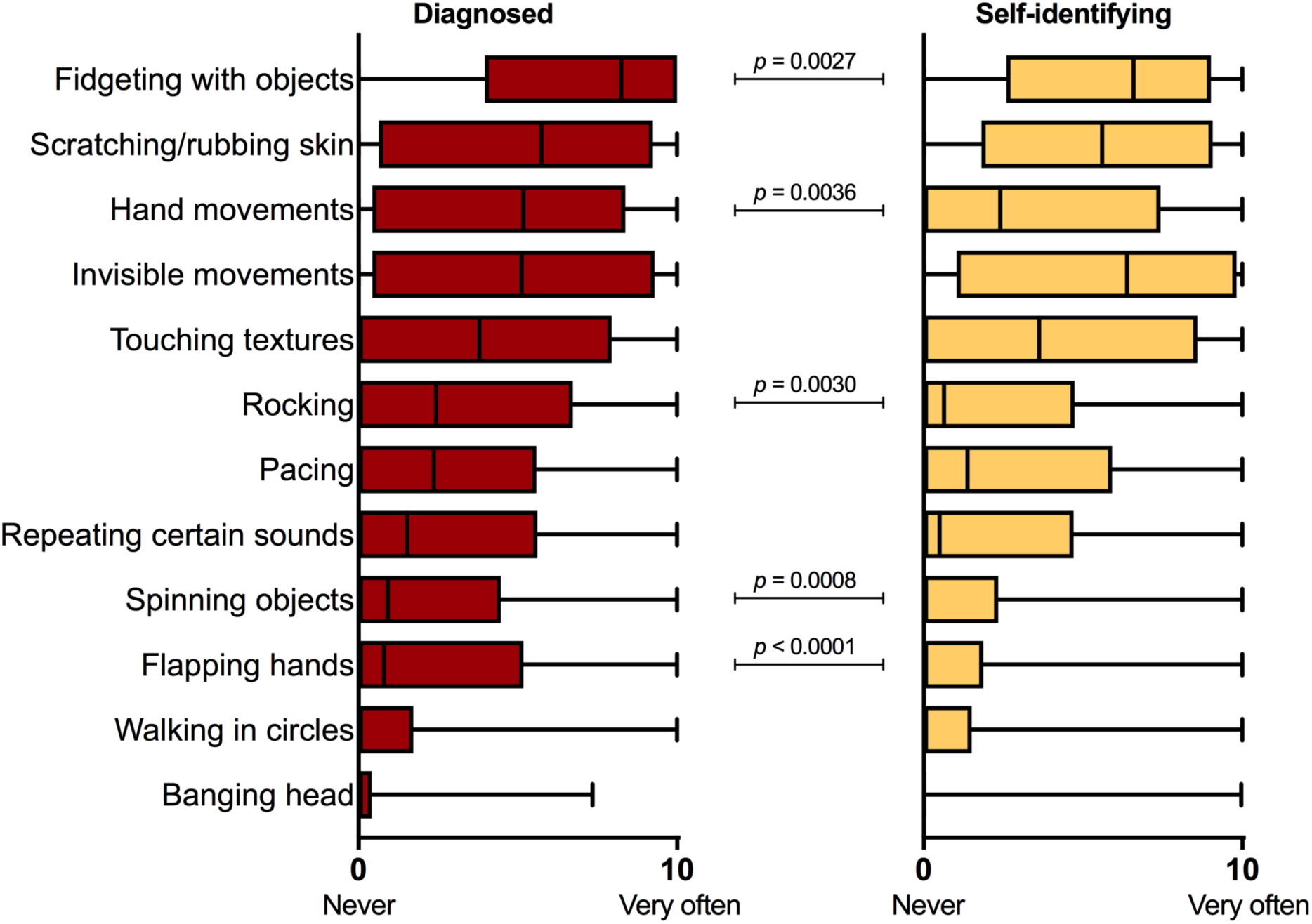
Self-ratings of selected behaviors. Shown are the interquartile ranges of the scores obtained in the diagnosed (left panel) and self-identifying (right panel) groups, presented in descending order based on the median score in the diagnosed group. All behaviors were seen in at least a proportion of participants (>25% with the exception of head banging in the self-identifying group). Higher scores in the diagnosed group were found for object fidgeting, repetitive hand movements, rocking, object spinning and hand flapping (Mann-Whitney tests with Bonferroni correction).

The most highly rated behaviors on our list were fidgeting with objects (median score 8.2 in the diagnosed group), scratching/rubbing skin (median score 5.7), repetitive hand movements (median score 5.2), invisible muscle activity (median score 5.1), and touching textures (median score 3.8). Rocking, pacing and repeating sounds seemed less common (median scores of 2.4, 2.4 and 1.5, respectively), and the median scores for spinning objects, hand flapping, walking in circles and head banging were low (range 0– 0.9). Mann-Whitney tests with Bonferroni correction indicated lower scores of fidgeting with and spinning objects, repetitive hand movements, hand flapping and rocking in self-identifying participants (**Fig. 4**). The between-group difference in head banging scores was associated with a low *p* value (uncorrected *p* = 0.0061), but did not pass Bonferroni correction.

The sum of all 12 scores was calculated for each participant as a “total RM score” (range 0–120), and cumulative distributions were compared between the diagnosed and self-identifying groups. The score itself does not mean anything specific, but reflects the degree of RMs reported by each participant. The majority of participants scored above 20, reflecting a combination of two or more behaviors (**Fig, 5A**). The between-group difference was not statistically significant (*p* = 0.087, Kolmogorov-Smirnov). The score was significantly correlated with the AQ score, but this relationship was variable on the subject level (Spearman correlation; **Fig. 5B**).

**Figure 5.**
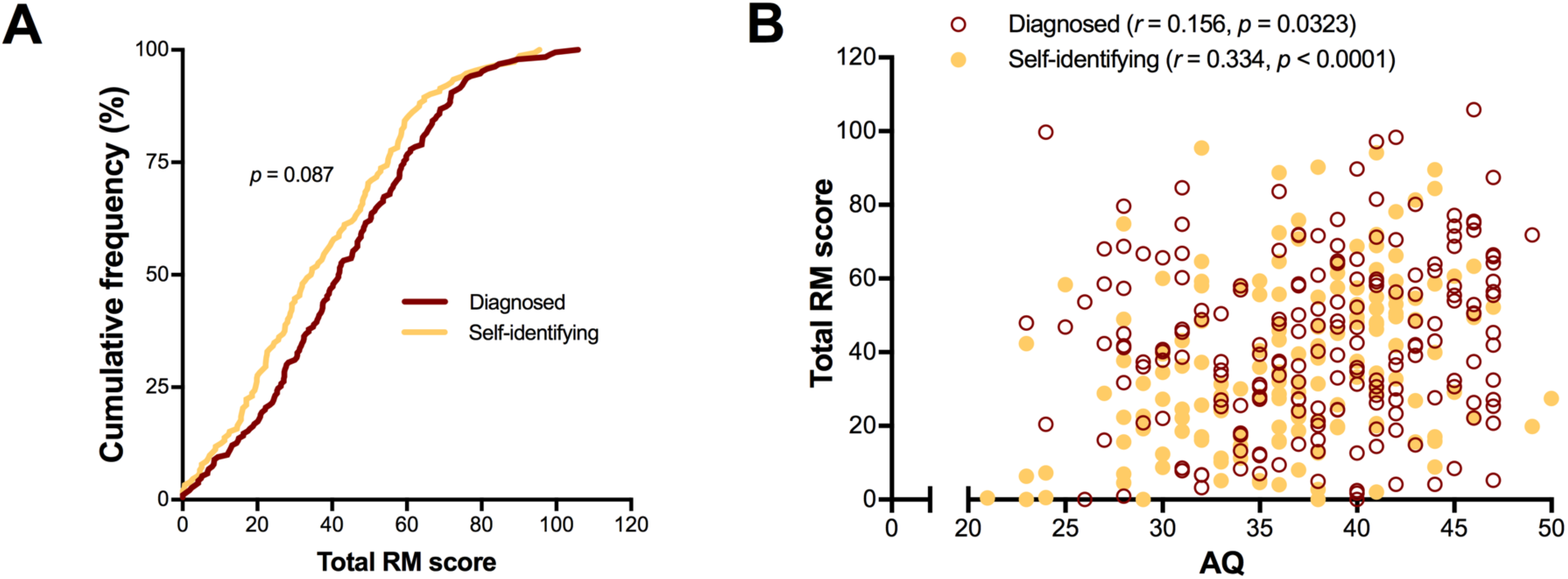
Distributions of total RM scores. A. Cumulative distributions of the total RM scores in diagnosed and self-identifying individuals, showing no significant difference between the groups. The majority of participants had a total score >20. B. Scatter plot of the total RM score against the AQ, showing a significant positive Spearman correlation, especially in the self-identifying group.

We asked whether the RMs had been expressed in childhood, and if so at which ages, but the responses were of very variable quality (the question was likely too broad given that most participants expressed a combination of several RMs). In the diagnosed group, 13% left the textbox blank, and in the self-identifying group, 16% did. Twelve percent of diagnosed and 9% of self-identifying participants either did not remember or provided responses that we could not interpret. A confirmatory response was seen in 75% of diagnosed and 74% of participants. Only 3 participants explicitly denied having expressed the RMs in childhood, and 2 of them expressed few RMs in adulthood (total RM scores 0, 4.2 and 28). There were multiple references to a lack of awareness before diagnosis or before self-monitoring increased with age. The results should be interpreted with caution, but suggest that the majority of participants had begun expressing RMs during development.

We asked participants to describe any RM habits that had not been covered by the list. This field was filled out by 62% of diagnosed and 64% of self-identifying participants. Non-motor RRBs that are also commonly referred to as “stimming” in the autistic community were sometimes reported in this textbox, including olfactory stimulation (smelling things), auditory stimulation (e.g. repeating the same song), visual self-stimulation (e.g. staring at Christmas lights), higher-order behaviors (e.g. organizing objects, counting, doing repetitive creative tasks), and vocal/verbal behaviors. We excluded non-RM behaviors as our question concerned motor stereotypies. A variety of RMs were described, and often in combination. The majority of them involved 1) upper extremities (19% and 22% in diagnosed and self-identifying, respectively), 2) lower extremities (19% and 25%), 3) mouth/throat (22% and 25%), 4) whole body/locomotion (15% and 11%), 5) picking/pulling of skin/hair (7% and 13%), and 6) playing with hair (twirling, twisting, touching, etc.) (7% and 11%). Among participants who described RMs involving upper extremities, the majority described finger movements or discreet hand movements (e.g. tapping/flicking fingers, rubbing/clasping/clenching hands, digging nails into skin). Among participants who reported lower extremity movements, the most common behaviors were also relatively subtle (e.g. jiggling/bouncing legs, wiggling/tapping toes or feet). Mouth/throat behaviors involved e.g. biting nails, biting self, chewing inside of cheek, chewing objects, clicking tongue or throat, sucking thumb, or clenching teeth. Whole-body/locomotive behaviors included jumping, bouncing, spinning, swaying, running, or tiptoe-walking.

### Camouflaging of repetitive movements

We next asked whether participants had tried to hide their RMs in childhood, adolescence and adulthood, giving the options of Always, Sometimes, Never, Don’t know and N/A.

We found that 80% of diagnosed participants and 78% of self-identifying participants had tried to hide the behaviors “sometimes” or “always” in adulthood, and 85% of diagnosed and 80% of self-identifying participants in adolescence (**Fig. 6**). About a fourth of the participants did not know or remember whether they had camouflaged in childhood, but 55% of diagnosed and 59% of self-identifying participants reported having tried to hide RMs as a child. Next, we asked how they had tried to hide the RMs, with intentionally broad phrasing to encourage free reflection. About half of the participants in each group described one or more strategies in more detail.

**Figure 6.**
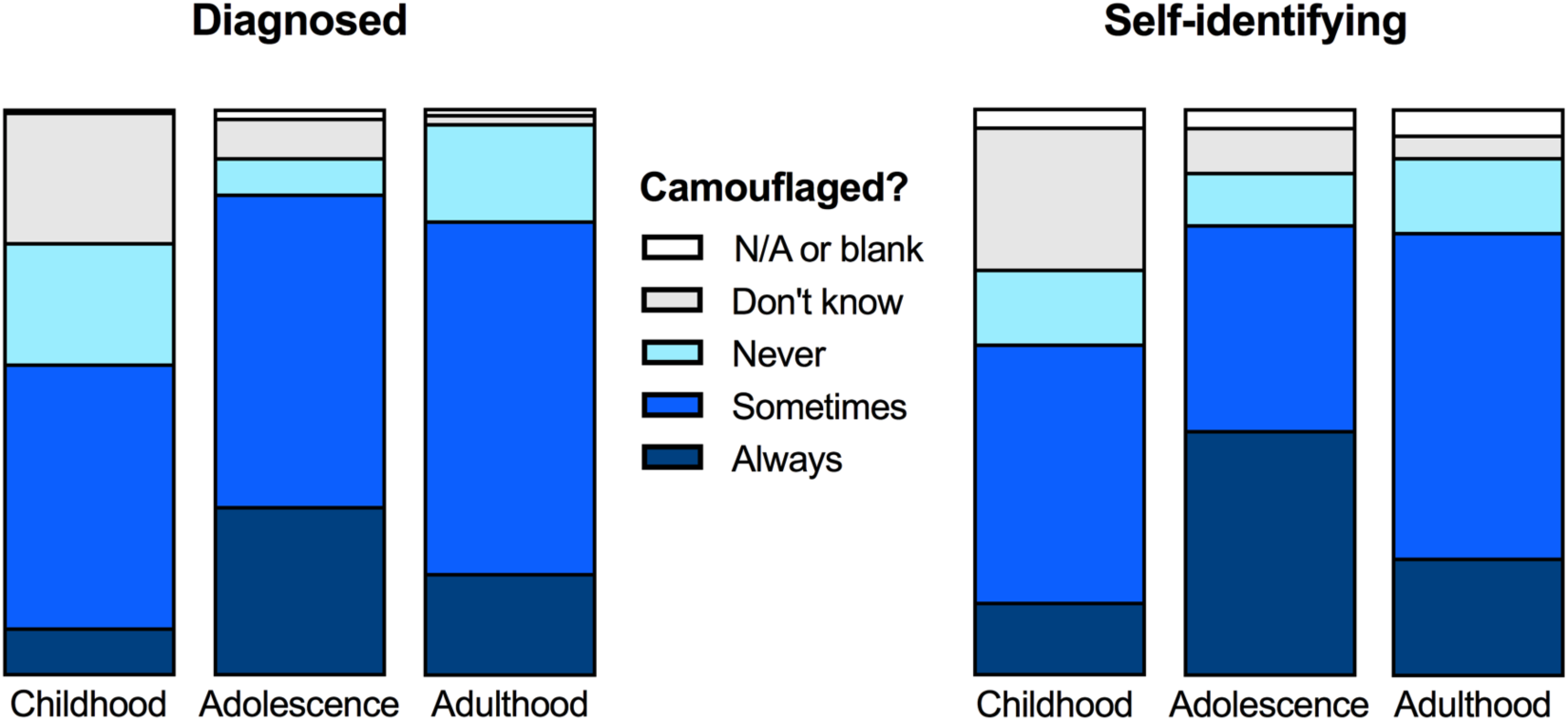
Camouflaging of RMs. Each bar illustrates the fractions of participants who had tried to hide RMs in childhood (left bar), adolescence (middle bar) and adulthood (right bar), with the two darker colors representing self-report of camouflaging (see legend in the middle). The majority of participants reported having camouflaged sometimes or always in all life phases.

We first analyzed the responses from the diagnosed group, but found that the same themes echoed through the self-identifying group. Common for all strategies was some type of self-inhibition. There were many references (47 participants) to having been bullied or disciplined for childhood RMs, and descriptions of various strategies to escape punishment. Five participants explicitly mentioned being physically punished (e.g. *“I used to get told off or slapped so I learnt to use things that weren’t obvious or were accepted /…/”*). Being verbally punished, discouraged from, or told off for RMs by parents or teachers was described by 25 participants (*e.g. “As a child I would get called out and/or punished. So I learned to be more discreet.”).* In addition, a few participants referred more indirectly to negative social reactions (e.g. *“I always had the urge to rock and stim, but recognized very early that it was considered weird or a sign of mental illness. I did it secretly when I was little and as I grew older stopped any obvious stimming.”* or *“/…/The negative reaction to any fidgeting or audible stimming was enough to make me suppress it.”).*

Participants frequently showed great awareness of the social consequences of their own behavior. Overall, it seemed common to want to be part of a social context, and to make great efforts to achieve this despite chronically feeling out of place. Eighteen participants explicitly mentioned feeling shame or embarrassment (e.g. *“I am aware of how weird my hand flapping looks. And the attention is embarrassing.”*), and another 34 participants mentioned in more or less direct ways that they had been self-conscious and tried to fit in socially (e.g. *“I knew it wasn’t normal or acceptable so I’d make my movements so small and [discreet] no one would notice.”*, or *“As a teenager I was hyperaware of my body language despite being clumsy – I wanted to avoid teasing and attention. I had no notion of being autistic, I just didn’t want to be ‘weird’.”*). A few participants described how they had learned very early in life to tense their muscles or dissociate in order to stop any muscle movement (e.g. *“/…/ I learned to sit completely still when required (I was able to sit through the ballet at the age of 5 without moving a muscle).”* or *“/…/ I was told that it made people think I was sick or broken, so I learned to dissociate from my body and sit very still./…/”*).

We saw three behaviorally separable strategies within the responses, though more than one strategy was often used within one participant. For the purpose of this report, we call the strategies “*Substitution*”, “*Self-isolation*” and “*Active suppression*”. Example quotes are shown in **Tables 1–3**. We do not make statistical comparisons as it is possible that a greater number of participants may have had responded if asked explicitly about the different strategies.

**Table 1.**
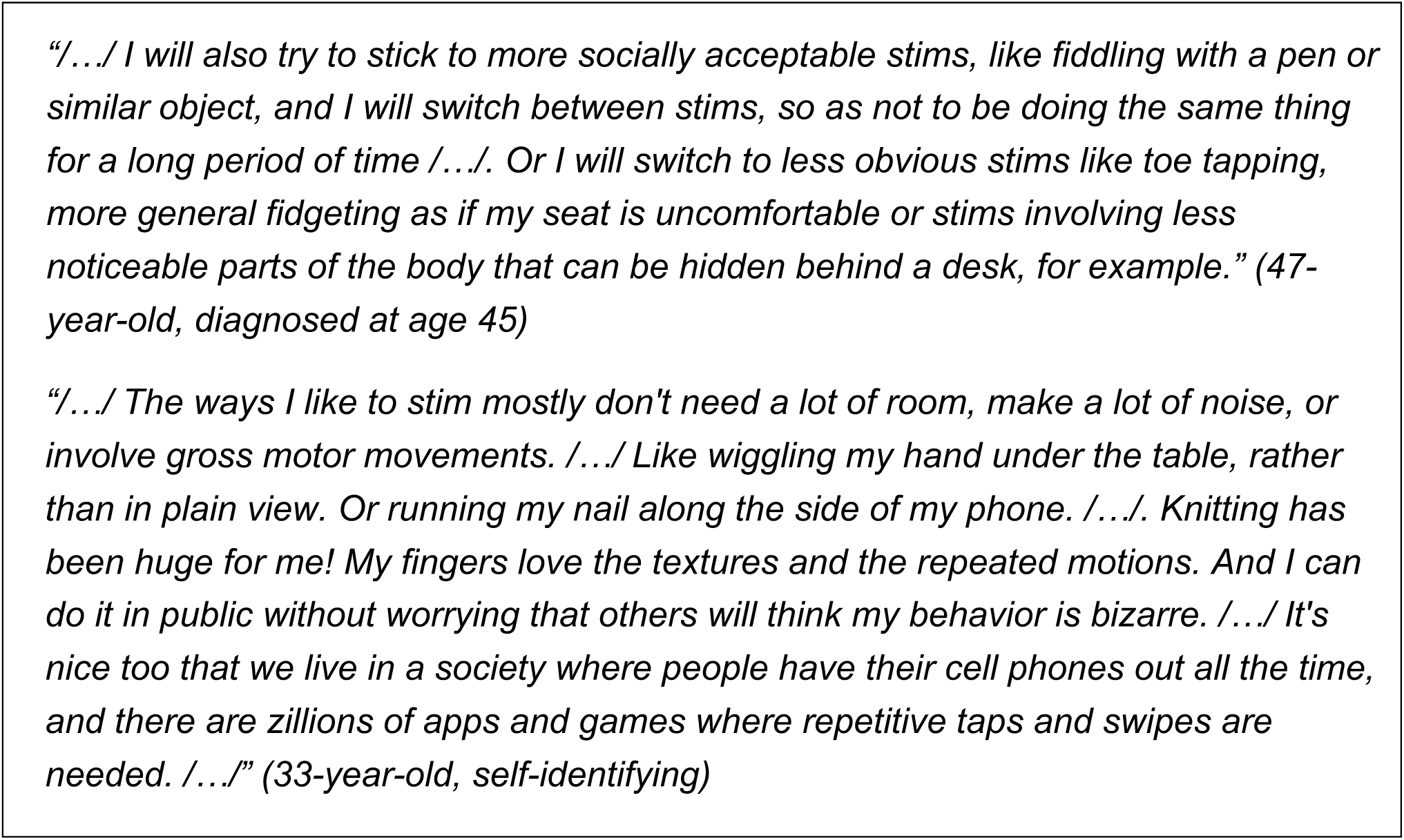
Example quotes: Substitution for a discreet or acceptable movement

With *Substitution*, we refer to the substitution of an obvious or stigmatizing behavior for something more socially acceptable or something more discreet. This was described by 73 diagnosed and 40 self-identifying participants. Behaviors mentioned here were ones that could be turned into something more discreet, like fidgeting with objects inside a pocket or under a table, biting inside of cheeks, moving toes inside shoes, or simply using very small movements. Socially acceptable expressions that were mentioned included fiddling with pens or other mundane objects, chewing gum, tapping feet, or walking in a circle while speaking on the phone. Several participants mentioned choosing behaviors that many other people have as nervous habits, because anxiety is less stigmatizing than autism-related RMs. Participants also sometimes mentioned having picked up more negative habits in order to hide their needs. For example, there were descriptions of taking up smoking to fulfill “oral sensory needs” or as a socially acceptable way to fidget with an object. Hidden self-injurious behaviors like chewing on lips or cheeks were described as a response to the social need to avoid more obvious RMs. One participant reported “self-medicating” with alcohol with the purpose of achieving a similar effect. Other participants mentioned using creative activities, like sewing, embroidery, knitting, or spinning yarn, as RMs that fulfilled their sensorimotor needs without being stigmatizing. (**Table 1**)

With *Self-isolation*, we refer to the isolation of behaviors to an environment where no one could judge or see. Participants described that they restricted their expressions of certain behaviors to home or a safe social environment, and generally included isolating themselves from other people. This was described by 53 diagnosed and 48 self-identifying participants. (**Table 2**)

**Table 2.**
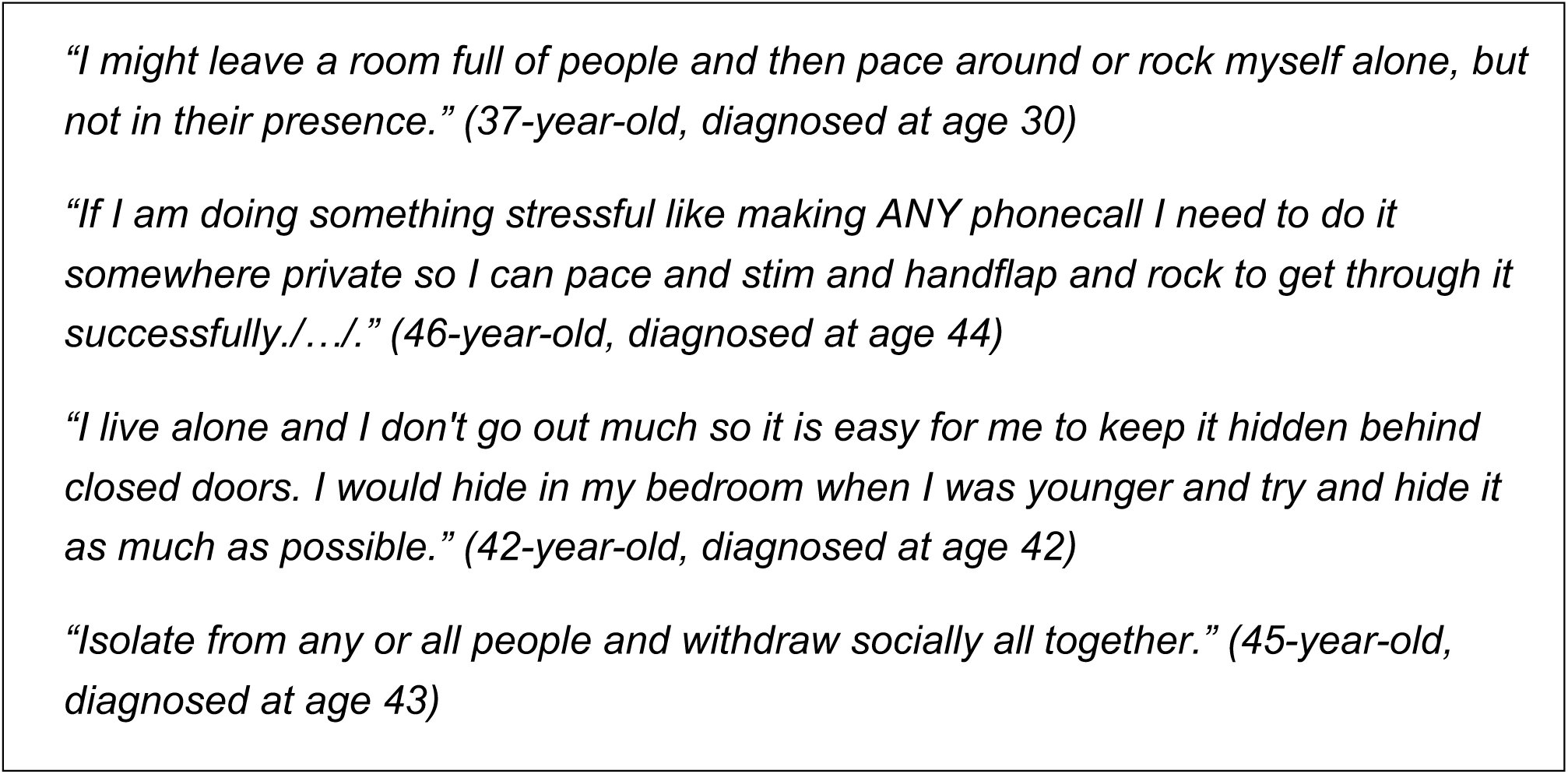
Example quotes: Self-isolation

Lastly, we coded a strategy as *Active suppression* when there was a description of a more forceful suppression of a behavior, without the replacement by another behavior (Substitution) or without removing oneself from others (Self-isolation). This was described by 27 diagnosed and 21 self-identifying participants. In this category, participants described an active effort to suppress the motor activity. This could be achieved by mental self-control or physical restraint (e.g. sitting on hands or making the hair inaccessible by braiding it). Active suppression was commonly described by participants who also utilized Substitution strategies in other contexts. (**Table 3**)

**Table 3.**
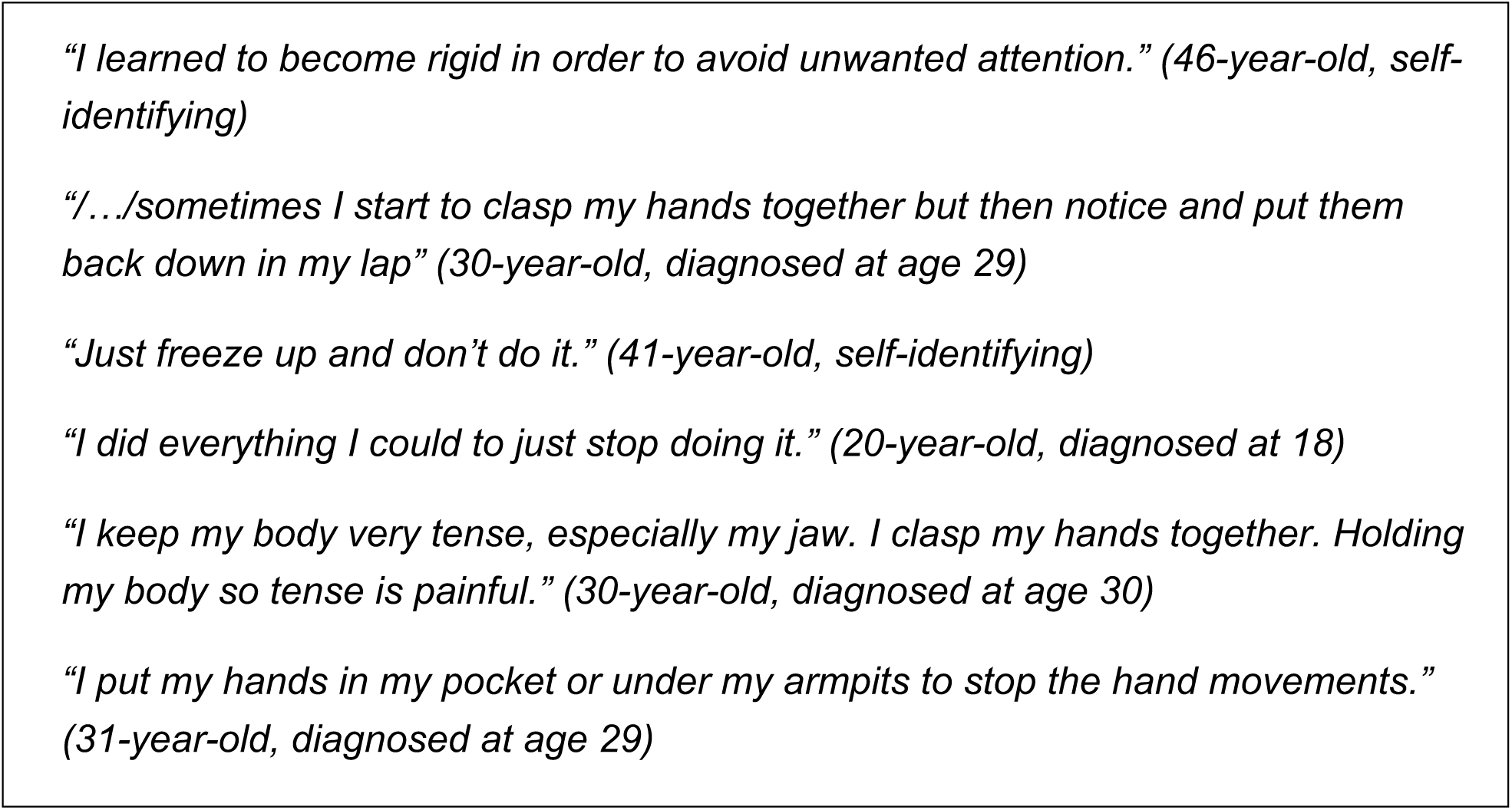
Example quotes: Active suppression

### Participants’ view on their need to camouflage better

Motor stereotypies are often viewed as an undesired symptom of autism, but many (though not all) participants in our cohort were opposed to this view, even though most still camouflaged their own RMs. When we asked if they wished that they could control their behaviors better, the majority of the diagnosed group said “No” (60%), 35% said “Yes”, and 5% responded “N/A”, “Don’t know” or left the question blank. Of those who answered “No”, 77% reported camouflaging in adulthood (sometimes or always), and of those who answered “Yes”, 88% did. The same pattern was seen in the self-identifying group.

Additional comments were provided by more than half of all participants. These comments indicated that the dominant reason for wanting better control was when behaviors had negative physical consequences (e.g. head impact, damage to skin, or pain from repetitive motion). The second most common reason was related to the fear of making a negative impression, draw attention, or cause discomfort in other people, at work and elsewhere. Among people who did not wish to control their RMs, the main topic of the comments was the opinion that these behaviors are harmless and help with self-regulation. It is probably reasonable to believe that some participants who responded “No” did so because they already felt they had sufficient control, but very few comments actually expressed that.

### Negative aspects of camouflaging

After noticing the abundance of spontaneous comments on negative aspects of trying to control RMs, and discussions about social acceptance and self-regulation, we looked more closely at these topics. We went through all text responses related to camouflaging and control of RMs to identify relevant comments. The summary below is derived from text from 76 respondents in the diagnosed group and 50 in the self-identifying group. Like in the previous analyses, the results from the different groups were virtually indistinguishable, so for this final section we pooled the participants (*N* = 126).

Comments on wanting acceptance from others were entered by 52 participants. About half were relatively simple comments like: *“I wish people could accept me if I did them, I enjoy them mostly so I wish it was OK to do in front of people.”*. Some expressed how they were actively trying to stop or had stopped camouflaging (33 participants) (*e.g. “I’m glad I have a word for these little habits now and am interested to learn more about healthy stimming”*, or *“Since I realised I was autistic, I have stopped stopping myself from rocking at home”*). Many viewed camouflaging as a maladaptive response to an unaccepting social environment (e.g. *“It’s quite difficult but as I grow older, I find that hiding them is more about others’ comfort rather than my own. I’ve been putting my comfort before [theirs] as I mature.”* or *“I try to stim without shame now, but hiding it has been hardwired into my brain so it’s difficult to get out of the habit.”*).

A number of participants advocated for viewing RMs as a normal part of the behavioral repertoire of many autistic people and argued that it would be most constructive to allow their expression (e.g. *“While it is useful to be able to choose to stim or not, I’m not sure it helps kids to try to force them to control it, or hide it. I hold a lot of tension in my body from trying to control myself all the time.”*). RMs were discussed as an important means of emotional self-regulation and, in a few cases, non-verbal communication. Comments along these lines often contained the disclaimer that self-injurious behaviors do need to be addressed (e.g. “*Unless a stim is self-injurious (or injurious to others), there’s nothing wrong with engaging in stimming. It’s a calming mechanism and efforts to quash it are misguided.”*). Participants described how their autism diagnosis or “self-diagnosis” had helped them understand their instinct to move in these ways, explore possible advantages with RMs, and identify and evaluate camouflaging behaviors (e.g. *“One of the reasons I wish I had been identified as autistic earlier is so that I could have had the benefit of stimming as and when needed to help self-regulate. I feel I have been repressing this natural instinct.”*).

## Discussion

This exploratory study shows that RMs do occur among cognitively and verbally able autistic adults, but are often hidden from public view through camouflaging. These findings mirror research in the social domain and in the domain of circumscribed interests, where non-male individuals show subtler or fewer diagnostic behaviors (Lord *et al.* 1982, Kopp and Gillberg 1992, Hartley and Sikora 2009, Kopp and Gillberg 2011, Mandy *et al.* 2012, Szatmari *et al.* 2012, Hiller *et al.* 2014, Lai and Baron-Cohen 2015). Camouflaging of RMs has to our knowledge not previously been reported, and may be important to consider to a greater extent in the clinical setting. Possible negative consequences of camouflaging should also be investigated further, so that this aspect can be considered in the design of early interventions and parent/teacher education.

We believe that this study in part reached the “lost generation” of autistic adults (Lai and Baron-Cohen 2015), many of whom appear to have turned to social media for support and kinship, sometimes after many disappointing encounters with clinicians and scientists. While on-site studies have a much better chance of phenotypic characterization and clinical assessment, they attract a different population. A large proportion of our participants would not participate in such studies. Some find the setting aversive and pathologizing, many lack the energy for travel or social interactions, and many have expressed to us how much easier it is for them to communicate in writing than in person. Thus, our sampling bias was different and in many ways complementary to the sampling bias of on-site studies.

We reached a large number of undiagnosed participants, including individuals awaiting the outcome of an evaluation, people who have not found a specialist nearby, and people whose primary care physicians do not agree that a referral is needed. The striking similarities between diagnosed and undiagnosed participants are consistent with a clinically relevant prevalence of autism in the undiagnosed group. It is important to identify barriers to evaluation or diagnosis for individuals who fly under the radar due to well-developed camouflaging strategies, and improve our understanding of the diverse manifestations of autism. Autism studies can only be done without sampling bias once all individuals who do meet criteria can get a diagnosis, and once those with a non-binary gender are included.

The significant representation of transgender participants should serve as encouragement to recruit transgender people in future studies. The inclusion of non-binary/transgender participants in our study allows for a first glance at this group, which is excluded from many studies through the binary definition of sex. The diversity of gender identities seen in this study, as well as the comments that were disapproving of the very question, illustrate the importance of aligning study terminology with the preferences of transgender communities, whilst keeping the question comprehensible to those cisgender individuals who still lack insight into gender diversity. Through confidential approaches from non-binary and transgender individuals during preliminary studies, we understood that it was unacceptable for them to just be asked about “biological sex”. By including enough options, we reached a large group of people who were very generous with their experiences of autism.

Even though camouflaging is currently a hot topic when it comes to autistic females, an unresolved and sometimes forgotten question is how common it is among cisgender males to differ in similar ways from the “classical” male phenotype. Such individuals may also represent a hidden and underdiagnosed group, and they were not included in this study. As mentioned above, we found few cisgender males in the social recruitment venues. Even though many autism groups online did include many males, they appeared to use social media in a less personal way and were more difficult to reach through support groups. This problem can probably be overcome in future studies by optimization of recruitment strategies.

The participation of individuals of different nationalities and the online format limited our ability to ascertain that robust diagnostic procedures had been used in all cases. The self-report format also makes it possible that some participants erroneously reported an official diagnosis. While the diagnostic status of participants would ideally be better characterized, we do believe that the majority of the diagnosed group did reflect diagnosed autistic individuals: There was no obvious incentive to be dishonest. We actively sought participants both with and without formal diagnosis and there was no financial incentive to do the study.

The view of many participants that suppression of RMs had been damaging for them may be an important finding. That there are costs to camouflaging has been discussed in popular media, and was recently shown to be related to an increased risk of suicidality (Cassidy *et al.* 2018). Participants in this study wrote that camouflaging of RMs during development had caused challenges in adulthood with self-acceptance and general functioning. In addition, there seemed to be an unconscious component to camouflaging – an automatic adaptation to social demands. This could be viewed as a natural part of development, but many of our participants felt that RMs were helpful with concentration, sensory function, or emotional processing. It could be speculated that RMs provide an input to the brain that compensates for some neurodevelopmental difference that causes difficulties in daily life. If that is the case, early interventions might need to be optimized to account for that.

In conclusion, the current study reached a large number of individuals within a rarely studied subpopulation of the autism spectrum. Many would be characterized as belonging to the most “high-functioning” population, given high levels of education and relatively high rates of employment. However, poor psychological well-being and delayed diagnoses indicate that this group is highly clinically relevant and needs more attention and better access to assessments. Our finding that RMs can often be self-monitored and camouflaged reveals a new important factor to consider in scientific and clinical settings.

## Acknowledgments

The study was supported in part by Autism Stiftelsen, The Autism and Asperger Association, Sweden, and by crowdfunding from anonymous donors on social media. We thank all the participants for putting in so much time and effort and for giving us feedback on everything we do. Our neurodiverse advisors have been very helpful in improving our communication with autistic/transgender individuals. We especially thank Katherine Lawrence and Cori Frazer for sharing their insights to help with terminology and study design. Thanks also to all members of the Extraordinary Brains outreach team for contributing to establishing our online presence.

